# Meta-analysis of the General Coral Stress Response: *Acropora* corals show opposing responses depending on stress intensity

**DOI:** 10.1101/817304

**Authors:** Groves Dixon, Evelyn Abbott, Mikhail Matz

## Abstract

As climate change progresses, reef-building corals must contend more often with suboptimal conditions, motivating a need to understand coral stress response. Here we test the hypothesis that there is a stereotyped transcriptional response that corals enact under any stressful conditions, functionally characterized by downregulation of growth and activation of cell death, response to reactive oxygen species, immunity, and protein homeostasis. We analyze RNA-seq and Tag-Seq data from 14 previously published studies and supplement them with four new experiments involving different stressors, totaling over 600 gene expression profiles from the genus *Acropora*. Contrary to expectations, we found not one, but two distinct types of response. The type A response was observed under all kinds of high-intensity stress, showed strong correlations between independent projects, and was functionally consistent with the hypothesized stereotyped response. Higher similarity of type A responses irrespective of stress type supports its role as the General Coral Stress Response providing a blanket solution to severely stressful conditions. The distinct type B response was observed under lower intensity stress and was weaker and more variable among studies than type A. Unexpectedly, the type B response was broadly opposite the type A response: biological processes up-regulated under type A response tended to be down-regulated under type B response, and vice versa. Gene network analysis indicated that type B response does not involve specific co-regulated gene groups and is simply the opposite of type A response. We speculate that these paradoxically opposing responses may result from an inherent negative association between stress response and cell proliferation.

## Introduction

Coral reefs provide disproportionately high ecological and economic benefits but are among the ecosystems most threatened by climate change. This has motivated efforts to understand mechanisms of coral resilience to environmental stress. A popular method for approaching such questions is profiling genome-wide gene expression responses to stressful conditions. Often, these studies focus on a single stressor and seek to identify genes and pathways that underlie the adaptive responses to it. However, in isolation, these studies cannot differentiate between gene regulation specific to a particular stressor, and gene expression changes reflecting stress in general. For this reason, delineation of corals’ general environmental stress response is needed to understand how they contend with suboptimal conditions.

The concept of a generalized stress response has been investigated thoroughly in prokaryotes. Here, *the general stress response* refers to a set of genes coordinately induced under diverse stressful conditions by the activity of alternative sigma factors, which competitively bind with RNA polymerase to preferentially transcribe stress response genes (Ron 2006). Conditions include starvation, acid stress, osmotic shock, and temperature shock among others. In *E. coli*, the general stress response is driven by alternative sigma factor σS, which directly or indirectly regulates up to 10% of the genome (Weber et al. 2005). The *E. coli* heat shock response is similarly regulated by alternative sigma factors σ^32^ and σ^E^ (Ron 2006). In the gram-positive bacterium *B. subtilis*, many of the heat shock proteins are regulated as part of the general stress response controlled by alternative sigma factor σ^B^ (Ron 2006).

General stress responses have also been described in eukaryotes. The transcription factor p53 has been described as a general stress response gene (Young et al. 2013), as it is activated not only by DNA damage, but additional stressors including hypoxia, oxidative stress, protein damage, and heavy metal toxicity (Abdulla and Campbell 1996; Hammond and Giaccia 2005). In *Sacharomyces cerevisiae*, Gasch et al. (2000) identified a stereotyped pattern of gene expression induced by diverse environmental stressors including temperature shock, nutrient limitation, oxidative stress, and osmotic shock, which they referred to as the *Environmental Stress Response* (ESR). The ESR was characterized by downregulation of growth-related processes, and upregulation of carbohydrate metabolism, detoxification, cell wall modification, protein folding and degradation, DNA damage repair, fatty acid metabolism, metabolite transport, vacuolar and mitochondrial functions, autophagy, and intracellular signaling. They proposed that this stereotyped expression pattern is a general adaptive response to suboptimal conditions (Gasch et al. 2000).

A transcriptional response resembling the ESR in yeast appears to exist in Cnidarians. Examining functional enrichment among heat stress genes in *Acropora hyacinthus* and in other coral studies, Barshis et al. (2013) pointed out conspicuous overlap with the yeast ESR (Gasch et al. 2000). Comparing heat stress responses among anemone strains, Cziesielski et al. (2018) describe a “core Cnidarian response to heat stress” including protein folding and oxidative stress genes. The idea of a general stress response was also described by Aguilar et al. (2019). Examining results from multiple RNA-seq studies, they point out that several complements of genes, including oxidative stress genes and HSPs, that are consistently identified in response to environmental stress. Indeed, there is a key set of gene functions mentioned in the majority of gene expression studies on coral stress. These include down-regulation of growth-related processes, and upregulation of misfolded protein maintenance, oxidative stress response, immune response, and cell death (Meyer et al. 2011; DeSalvo et al. 2012; Kenkel et al. 2013; Maor-Landaw et al. 2014; Table 1). Hence the existence of a general coral stress response has been widely hypothesized but has yet to be formally examined across diverse environmental stressors.

**Table 1:**
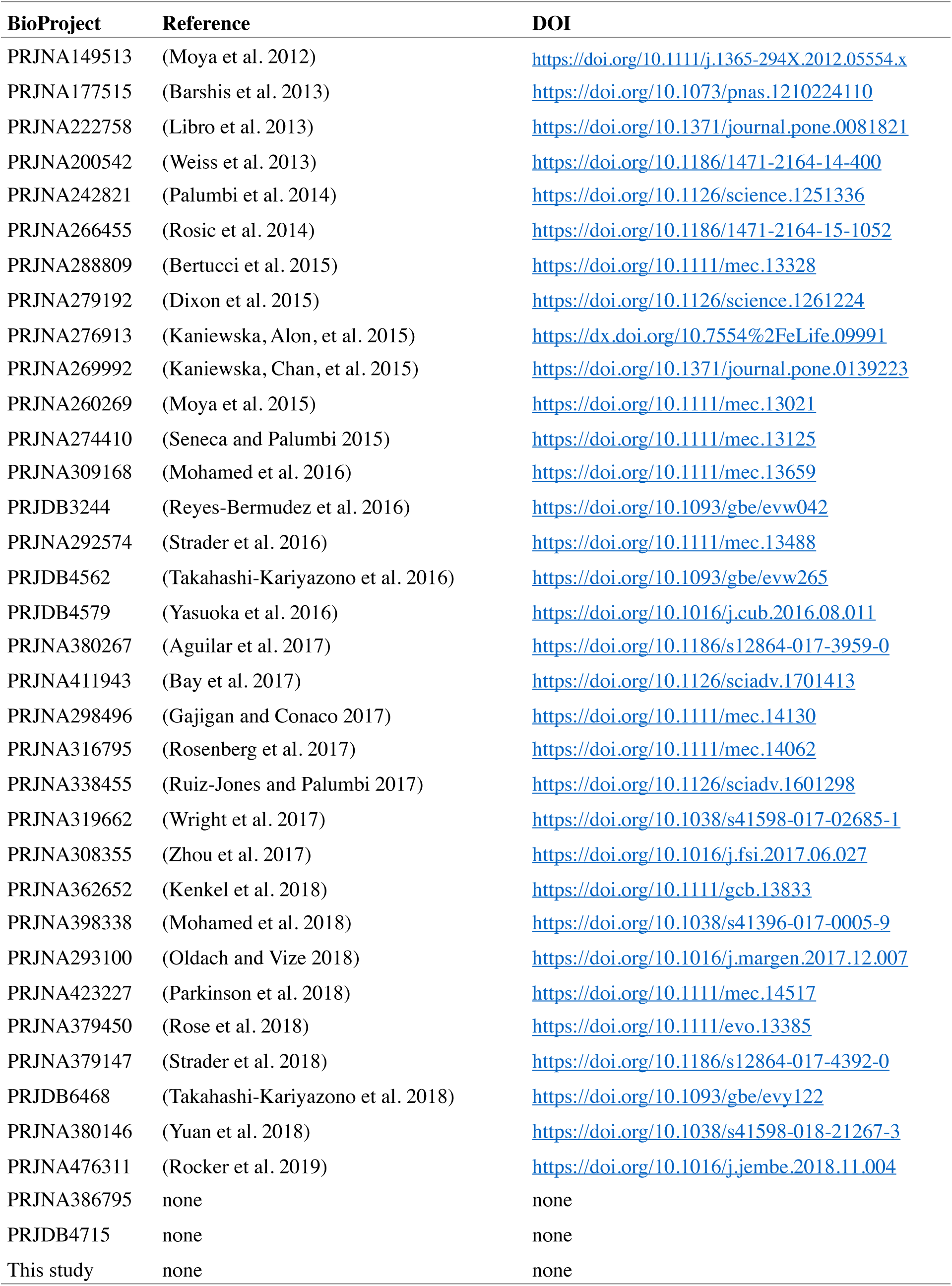
Data sources

Here we investigate the general stress response of reef-building corals. Specifically, we assess the hypothesis that there exists a core response that corals enact under any form of environmental stress. This stereotyped stress response can be contrasted with specific responses, that are induced only by specific stressors. The existence of a general stress response makes several predictions about coral gene expression under diverse stressful conditions: (1) gene expression responses to different stressors should be correlated, (2) gene expression responses to diverse stressors should demonstrate shared enrichment for the functions above, (3) it should be possible to identify sets of core genes up- and down-regulated by all types of stress. To test these predictions, we analyzed all publicly available RNA-seq datasets conducted on the coral genus *Acropora*, along with one new dataset.

## Methods

### Data sources

Relevant studies were identified by searching for the genus name *Acropora* on the NCBI SRA database. Only studies with Illumina data were used, giving 36 in total (Table 1). To make the datasets fully comparable, we obtained raw sequencing reads and mapped them all to the same reference, the recently assembled high-quality genome of *Acropora millepora* (available from https://przeworskilab.com/acropora-millepora-genome/).

The original SRA Run Table used for the study is shown in Table S1. Whenever possible, sample traits were filled in from information included in the SRA Run Table. When not possible, sample information was gleaned from the BioSample accessions, publications, and in a few cases, by directly asking the authors. The final modified version of the sample trait table used for analysis is saved as Table S2. The analysis also included RNA-seq reads from a new study in our laboratory.

Experiments for the new dataset included in this study were conducted at Orpheus Island Research Station in November 2018, under GBRMPA permit G18/41245.1. Four adult colonies of *Acropora millepora* were collected by SCUBA: three from Northeast Orpheus (labeled N1, N2, and N10), and one from Little Pioneer Bay (labeled L1), and transported back to raceways of unfiltered seawater. Nubbins were broken off from the colonies and maintained in the raceway on dish racks for 10-17 days before experiments were conducted. Treatments were intended to cause bleaching by acute stress.

### Cold experiment

The cold experiment included a cold and a control treatment group. Each treatment group comprised three replicate nubbins from each of the four colonies (N=12 nubbins per treatment group). Nubbins from each treatment group were hung from a fishing line in a 10 L container during the experiment. The cold treatment group container was placed in a refrigerator at 4°C for six hours, after which it was returned to ambient temperature. This treatment produced a ramp downward to 10.6°C over six hours followed by a ramp back to ambient temperature over another six hours (Figure S1). At the same time, the control group container was moved into a cabinet to mimic the dark of the cold treatment. Tissue samples were fixed in ethanol 17 hours after initiation of the 4°C treatment and immediately stored at −80°C. At this time point the corals showed visible signs of bleaching (Figure S1)

### Hyposalinity experiment

The hyposalinity experiment included a hyposalinity and a control treatment group. Each treatment group comprised three replicate nubbins from each of the four colonies (N=12 nubbins per treatment group). Treatments were applied in three replicate 5 L containers, each containing a single nubbin from the four colonies. The hyposalinity containers were filled with 42.8% filtered seawater (~15 ppt) by mixing 2.14 L of seawater with 2.86 L freshwater. We chose this treatment instead of less severe hyposalinity treatments because it caused bleaching in 100% of samples in trial experiments. Corals were moved directly into hyposalinity or control conditions from the raceway, and held with air circulation for six hours, after which flow of filtered seawater was resumed. Samples were fixed in ethanol 6 hours after beginning the exposure (immediately before resuming seawater flow). These samples already showed visible signs of bleaching. An additional set of samples was fixed 14 hours after beginning the exposure (8 hours after seawater flow was resumed). Samples were stored at −80°C immediately after fixing. A schematic of the experiment and example photos of control and treated samples are shown in (Figure S2).

### Heat experiment

The heat experiment included a heated and control treatment group, each with three replicate nubbins from each of the four colonies (N=12 nubbins per treatment group). Treatments were applied in single 10 L containers. The heated group was ramped to 36°C over three hours and held at 36°C for three additional hours before ramping back to 28°C over 3 hours. We chose this temperature to ensure bleaching in 100% of samples based on trial experiments using 34, 35 and 36°C. Samples were fixed in ethanol 14 hours after the treatment was started (5 hours after the returning to ambient temperature). Samples showed visible signs of bleaching at this point. For one of the colonies (N1), all three replicates had died by the sampling point. A second set of tissue samples was fixed in ethanol 25 hours after treatment was started. Samples were stored at −80°C immediately after fixing. Temperature traces for the experiment and example photos of control and treated samples are shown in (Figure S3).

### Multi-stress experiment

A final experiment was conducted with four treatment groups. These included a control group, a second heat treatment group, and two groups with multiple stressors. The heat treatment group was ramped to 35°C over three hours and held for three additional hours before ramping back down to ambient temperature (~28°C) over 2 hours (Figure S4). The multi-stress groups were exposed to a combination of hot and then cold temperature or cold and then hot temperature simultaneously with mild hyposalinity (71.4% filtered seawater; approximately 25 ppt). The hot-then-cold treatment group was ramped to 35°C over three hours, and then moved to a refrigerator at 4°C for three hours, which dropped the temperature to a low point of 24.2°C, then returned to an outdoor tabletop, where the temperature returned to 28°C over three hours (Figure S5). The cold-then-hot treatment group received the reverse; first placed in the refrigerator at 4°C for three hours, reducing the temperature to a low point of 19.4°C, then ramping to 35°C over a three hour period before it was moved to an outdoor tabletop allowing a return to ambient temperature over three hours (Figure S5). Tissue samples were fixed in ethanol 13 and 23 hours after the experiment began. Flow of filtered seawater was resumed after the first set of samples was fixed. By mistake, flow was not returned to the hot-then-cold treatment group. For this reason, this group was not fixed at the second time point. At 7.5 hours after the experiment began an additional set of tissue samples was fixed from a single colony (N10) because they appeared to be dying in the multi-stress treatment groups. These were taken in addition to the other tissue fixing time points.

### Library preparation

RNA was isolated using RNAqueous™ Total RNA Isolation Kits. Frozen coral tissue was submerged in lysis buffer and pulverized using a single 6mm diameter chrome steel bead (Biospec Cat No. 11079635c) and a Biospec Mini-Beadbeater-96 (cat. No. 1001) using 20-second beating duration. The resulting lysate was processed according to the RNAqueous protocol. Preparation of tag-seq libraries was carried out as described in (Kenkel and Matz 2016).

### Sequencing data processing

Detailed steps for the data processing pipeline, along with all custom scripts used during data processing are available on Github: https://github.com/grovesdixon/Acropora_gene_expression_meta.git. Fastq files for each previously published sequencing run were downloaded using the SRA toolkit. Adapter trimming was performed in paired- or single-end mode as appropriate for each study using cutadapt (Martin 2011), with a minimum length cutoff of 20bp and a PHRED quality cutoff set to 20. Quality of reads was assessed before and after trimming on a subset of 10,000 reads from each Run using FastQC (Andrews 2010). Reads were mapped to an annotated draft reference genome for *Acorpora millepora* that can currently available on the Przeworski laboratory website (Fuller et al. 2018). The reference genome was generated using a combination of PacBio reads and Illumina paired-end reads with 10X Chromium barcodes and anchored into chromosomes with linkage mapping data from two previous studies (Wang et al. 2009; Dixon et al. 2015). Mapping was performed with Bowtie2 in paired- or single-end mode as appropriate using the --local argument (Langmead and Salzberg 2012). Following alignment, PCR duplicates were removed using MarkDuplicates from Picard Tools (Broad Institute 2019). Sorting and conversion from sam files was performed using Samtools (Li et al. 2009). The reads mapping to annotated gene boundaries were counted using FeatureCounts (Liao et al. 2014). Mean absolute and relative read counts for each Bioproject are given in (Figure S6) and total read counts for individual samples in (Table S3).

### Principal component analysis

Overall variation in gene expression was assessed using principal component analysis (PCA). Raw read counts were normalized using the vst() function in the R package DESeq2 (Love et al. 2014) and the top 10% of genes with the greatest variance were used as input. With the full dataset, the first principal component correlated closely with raw read counts (R^2^=0.61). Based on this, we used the removeBatchEffect() function in R package limma (Ritchie et al. 2015) to control for the total number of reads counted on genes by FeatureCounts before PCA. The resulting PCA plot was assessed visually for clustering of samples based on BioProject (the original study the sequencing run came from), treatment, and developmental stage of the sample (gamete, embryo, larva, or adult). After controlling for read counts, the first principal component was found to correlate closely with developmental stage. Controlling for BioProject using limma::removeBatchEffect removed this correlation, ostensibly because each project typically had samples of just one life cycle stage, and larval samples were relatively rare. To preserve life cycle stage information, we present the PCA of the full dataset having only controlled for read counts. PCA was also performed on subsets of the data based on experimental treatments. Here we found a strong influence of BioProject, but less so for raw read counts. Based on this, for subsets of the data, we controlled for BioProject using limma::removeBatchEffect.

### Differential gene expression analysis

To identify genes differentially expressed in response to different stress treatments, we used DESeq2 (Love et al. 2014). A total of 15 projects were found to include stress treatment (Figure 2A). For all differential expression analyses, stress treatment was encoded as a binary variable, either ‘stress’ or ‘control’. For studies with intermediate stress treatments, for instance, a project that exposed corals to 350, 750, and 1000 ppt CO_2_, both the intermediate and extreme stress treatment were coded as stress. All tests included stress treatment and BioProject as predictive variables. For each dataset, genes with mean read count less than 3 were removed before differential expression analysis. Significance of stress treatment was tested using Wald tests.

### Discriminant analysis of principal components

Discriminant Analysis of Principal Components (DAPC) was implemented using the R package adegenet (Jombart et al. 2010). The function dapc() was run on the transposed matrix of variance stabilized read counts produced with DESeq2, retaining the number of principal components sufficient to account for 80% of total variance.

### Predicting stress treatment from gene expression

Logistic regression was used to build predictive models of stress treatment based on gene expression. Stress treatment for each sample was coded as the outcome variable, with normalized counts for each gene used as predictors after controlling for BioProject. Logistic regression using the lasso method was performed with the R package glmnet (Friedman et al. 2009). The value for lambda was selected based on the *lambda.1se* value returned by the *cv.glmnet* function, which is the largest value for lambda such that the cross-validated error was within one standard error of the minimum.

The Random Forest algorithm was also used to predict stress treatment from gene expression. As with logistic regression, predictors were normalized counts for each gene after controlling for BioProject. The algorithm was implemented using the R package randomForest (Liaw and Wiener 2002). The number of trees grown per iteration (*ntree* argument) was set to 500, and the number of variables (genes) randomly sampled at each split (*mtry* argument) was set to 1000. Validation of both logistic regression and Random Forest models was assessed using confusion matrices built with the R package caret (Kuhn 2019).

### Weighted gene co-expression network analysis

To identify clusters of coregulated genes associated with stress treatments we used Weighted gene co-expression network analysis (WGCNA)(Langfelder and Horvath 2008). Input for this analysis was the matrix of variance stabilized counts from the entire dataset after controlling for Bioproject using limma. Before network construction, we subset this matrix for the genes that fell within the top 75% for expression level and variance across samples. The resulting subset included 11284 genes (54.7% of total). We ran WGCNA with a soft threshold power of 12, a minimum module size of 30, and a module merging threshold of 0.3.

### Gene annotations

Gene annotations were acquired using eggNOG-mapper (http://eggnogdb.embl.de/#/app/emapper; Huerta-Cepas et al. 2017; Huerta-Cepas et al. 2019). This produced GO, COG, and KEGG and gene names (eggnog annotations). The annotation table is available as a (Table S4).

### Analysis of functional enrichment

Functional enrichment was assessed using two-tailed Mann-Whitney U tests as in (Wright et al. 2015), using package GO_MWU (https://github.com/z0on/GO_MWU). For differential expression between stress-treated and control corals, we used −log_10_ transformed p-values output by DESeq2, multiplied by −1 when the gene was down-regulated under stress, as input for the Mann-Whitney U tests. The test compares ranks among these transformed p-values to assess whether the distribution of ranks of genes included in a GO term diverges significantly from that of all other genes. The deviation of a GO term from the rest of the dataset can be quantified by its delta rank (the difference in the mean rank for the GO term and the mean rank for all other genes). Positive delta ranks indicate the GO term tends toward upregulation. Negative values indicate the GO term tends toward downregulation. The similarity of functional responses between datasets was compared by plotting the delta ranks against one another (Figure 4). It is important to note that these plots do not represent a formal statistical test, as the data points (Gene Ontology Categories) are not independent since they often encompass overlapping sets of genes, but are a convenient way of observing functional similarity or dissimilarity between two differential expression datasets. Functional enrichment for WGCNA modules was tested using Fisher’s exact tests on binary calls for module membership.

KOG (euKaryotic Orthologous Groups) analysis was performed using R package KOGMWU as in (Dixon 2015). This method is analogous to GO_MWU but uses only 23 non-overlapping functional classes of genes, which facilitates formal testing for functional similarity and concurrent visualization of multiple gene expression studies. To further contrast the different response types detected, we also performed KOG analysis on data from the yeast *Lachancea kluyveri* from (PRJEB10946; Brion et al. 2016). These reads were mapped to *Lachancea kluyveri* coding sequence reference acquired from Genome Resources for Yeast Chromosomes (http://gryc.inra.fr/index.php?page=home). Annotations for these coding sequences were acquired using eggNOG-mapper as described above.

## Results

Overall, developmental stage was the dominant biological source of transcriptional variation. After controlling for read counts, the first two principal components revealed clear clustering by Bioproject (the study that published the reads) and developmental stage (Figure 1A-B). The first principal component, which accounted for 22% of the variance, correlated with developmental stage, with recruits intermediate to larvae and adults (Figure 1B-C).

**Figure 1:**
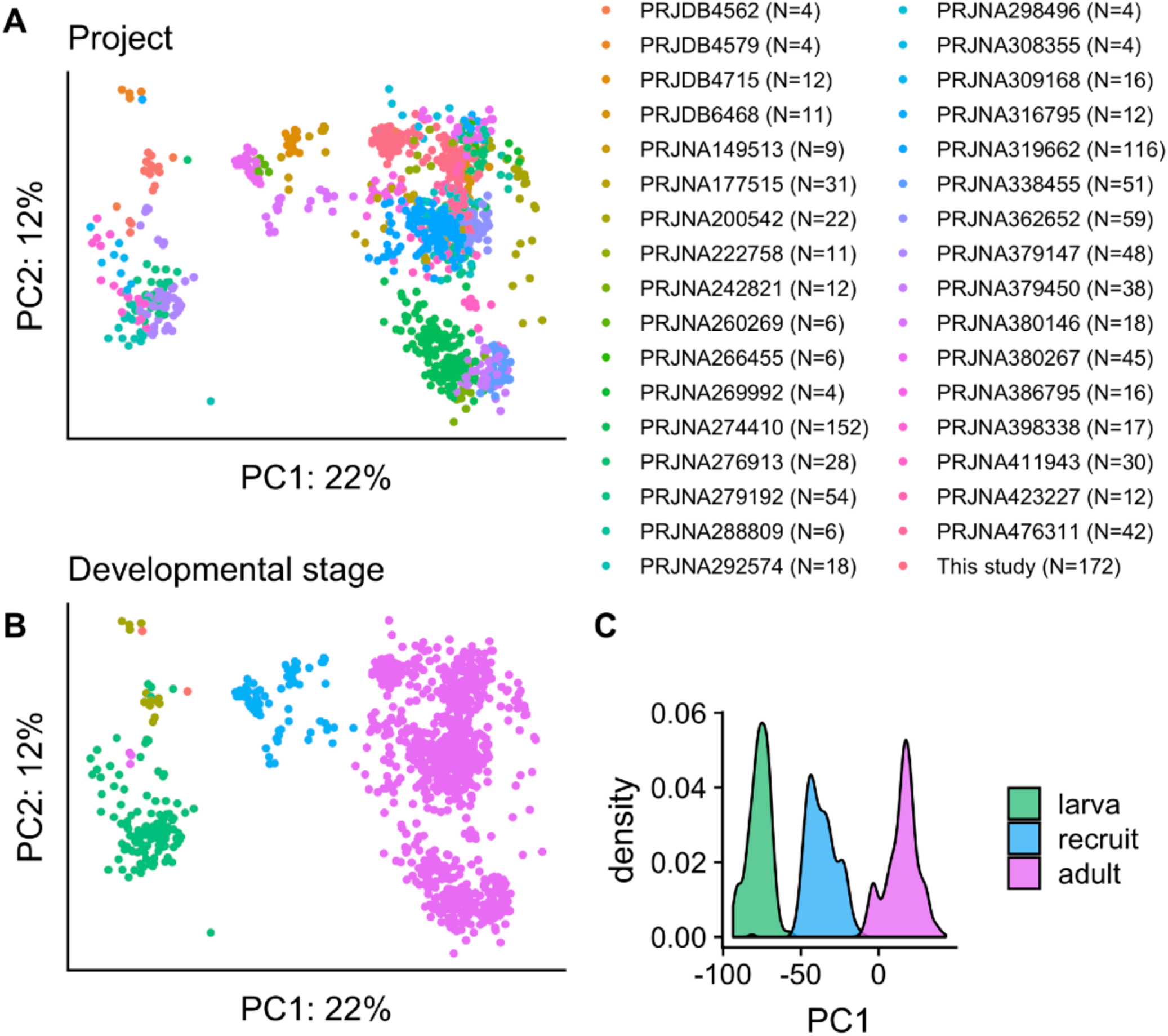
Principal component analysis (PCA) of full dataset. PCA was conducted on normalized reads counts with additional control for total reads counted per sample (see methods). (A) PCA color coded by BioProject allocation (the source of the published reads; 36 total projects). (B) The same PCA color coded by developmental stage (C) Density plot of PC1 for three primary developmental timepoints.

Counting this study, there were 15 BioProjects that included stress treatment, with 283 control samples and 335 stressed samples. These encompassed 6 types of stress: *heat*, *cold*, *hyposalinity*, *immune challenge*, *low pH*, and *multiple stressors* (Figure 2A). Many of the *heat*, *cold*, *hyposalinity*, and *multiple* samples were also experiencing bleaching (Figure 2A). Differential expression analysis comparing stressed to control samples across this full dataset (all stressed vs all controls) detected substantial differential expression, with more than half the genes significant at FDR < 0.1 (Figure 2B). After controlling for BioProject, PCA revealed that stress treatment was indeed a dominant source of transcriptional variation (Figure 2C). Discriminant analysis of principal components (DAPC), implemented to discriminate between stressed and control samples, nearly completely separated the stressed and control groups (Figure 2D) and correlated closely with the first principal component (Figure 2E). Based on the first principal component and discriminant axis, the *heat*, *cold*, and *hyposalinity* treatments tended to produce more severe transcriptomic responses than the *pH* and *immune challenge* treatments (Figure 2E). Notably, these were also the treatments that induced bleaching.

**Figure 2:**
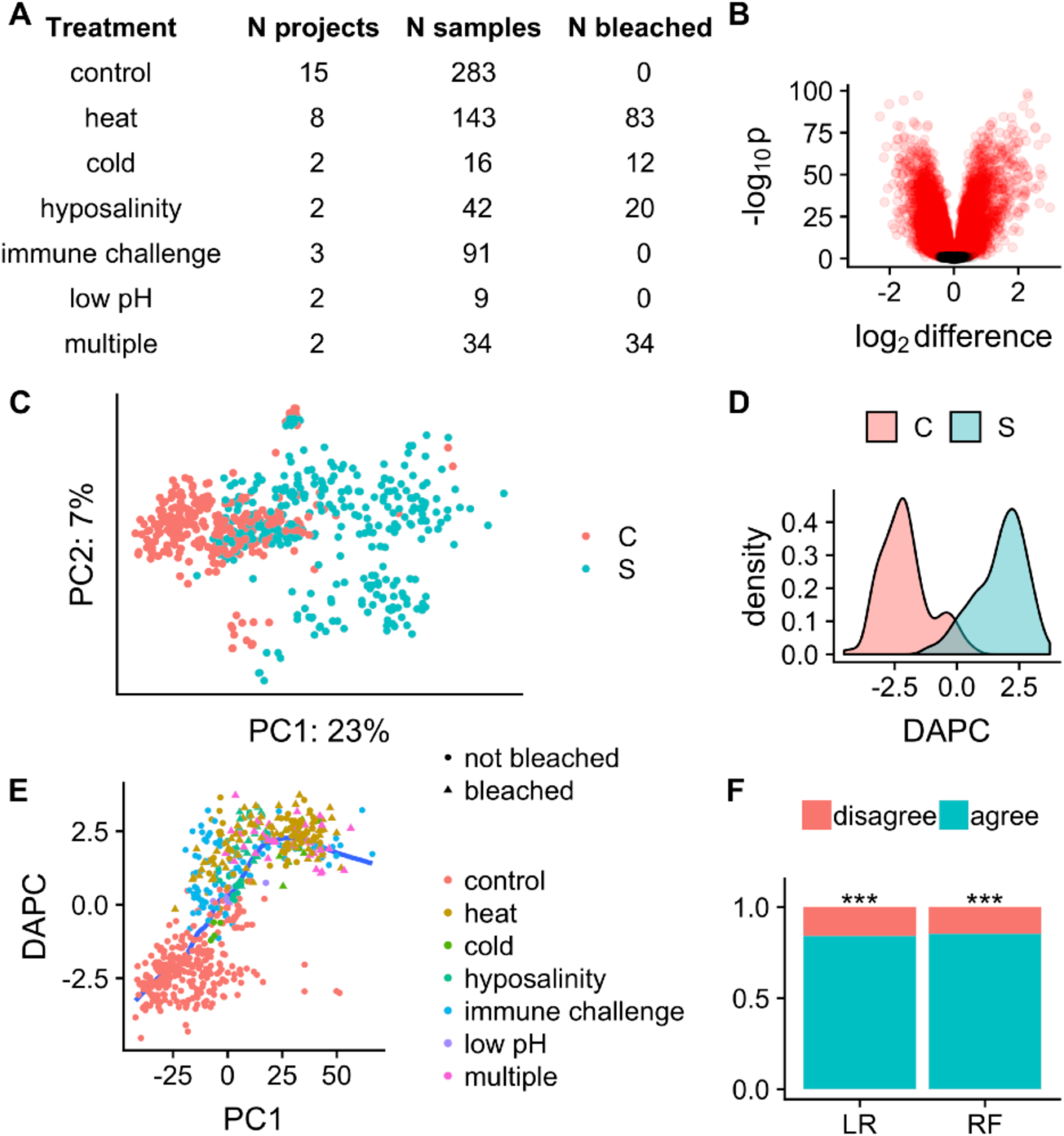
Transcriptional response to stress treatments. (A) Table showing the number of projects and samples for each type of stress. Analysis of these samples were performed coding samples as either stressed or control. (B) Volcano plot showing log_2_ fold differences and p-values for RNAseq counts between stressed and control samples. Positive log2 fold differences indicate upregulation in stressed samples. (C) Principal component analysis (PCA) of samples from all projects including a stress treatment. (D) Density plot of sample loading values for discriminant analysis of principal components (DAPC) performed to discriminate stressed and control samples. (E) Scatterplot of PC1 and DAPC loading values. (F) Barplot of accuracy percentages when predicting stressed status based on gene expression using logistic regression (LR) and random forest (RF). Models were trained on a random subset of 60% of the samples stratified by BioProject and accuracy was measured based on predictions made for the remaining 40% of samples.

We next developed classification models of a stressed transcriptome. Using lasso logistic regression and the random forest algorithm, we trained models to predict stress treatment from gene expression data using 60% of the dataset randomly sampled with stratification by BioProject. When applied to the withheld 40% of the data, both models predicted stress treatment with a minimum of 83% accuracy (Figure 2F). Together, these results indicate a similar transcriptional response in most stress-treated samples.

We next performed DESeq2 analysis for each of 14 published studies that included stress treatment and the four different experiments conducted for this study and compared the log_2_ fold changes from each of these to assess the similarity of their transcriptional responses. This comparison revealed a more complex situation than we hypothesized, with two distinct classes of response. Hierarchical clustering revealed that eight published projects, along with the four experiments from this study (88% of total samples), formed a tightly correlated group, with a mean Pearson correlation of 0.41 (Figure 3A). We refer to this group as type A response. Notably, the Bioprojects in this cluster included all stress types; hence their correlation is consistent with a general stress response paradigm. Surprisingly, a second group of six projects formed a distinct cluster of responses that were generally negatively correlated with the type A response, which we refer to as the type B response. Correlation of type B Bioprojects to each other was much weaker than for type A (mean Pearson correlation of 0.045). Although this may be due to lower samples size, repetition of the random forest model revealed the type B response was less predictable than the type A response, with type B samples frequently misidentified as controls (Figure 3B). Type B samples also accounted for a large amount of overlap between stressed and control samples on the first principal component and the discriminant axis (Figure 2C-E; Figure 3C-D), indicating less overall gene expression change among these projects.

**Figure 3:**
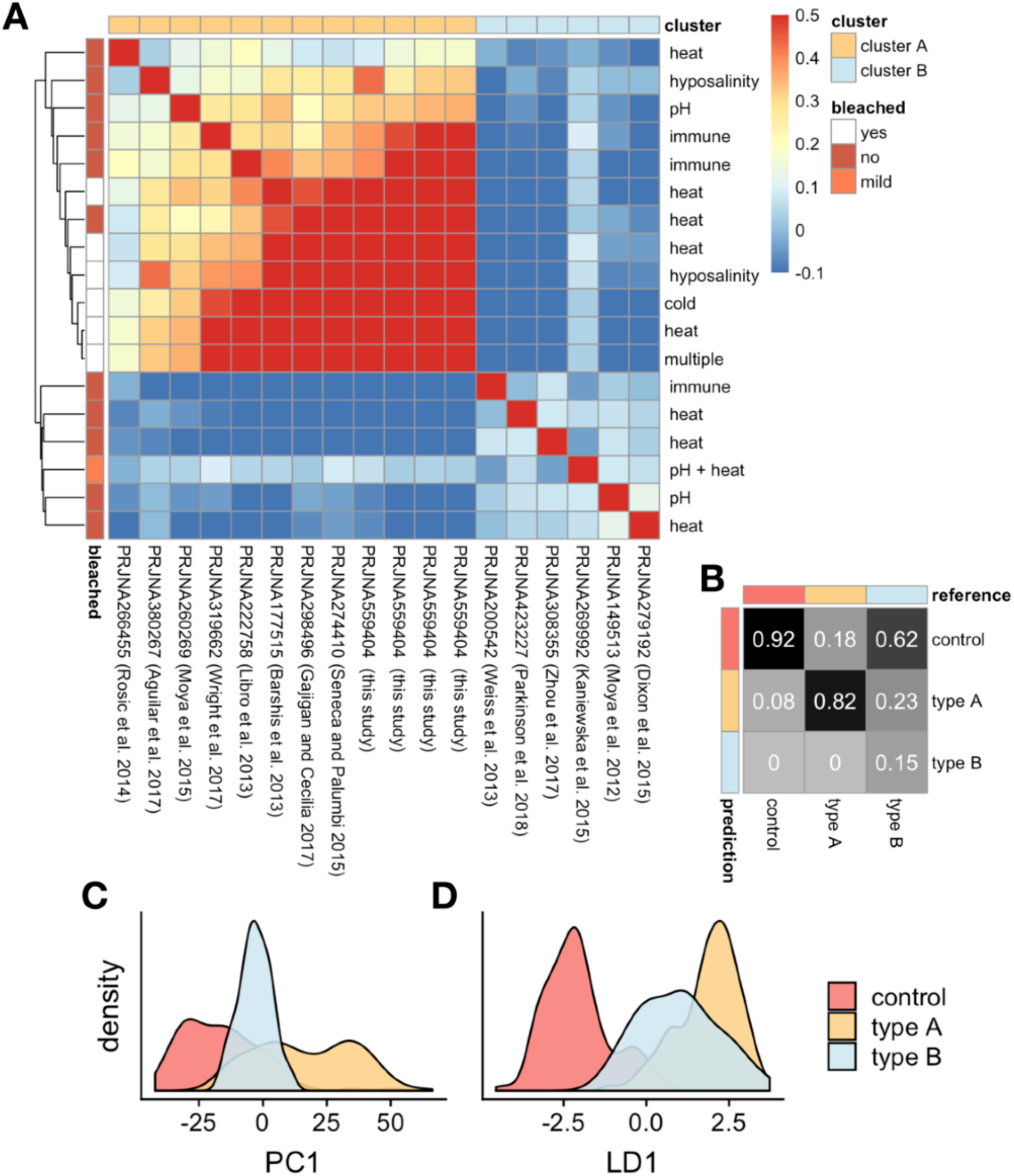
Identification of two clusters of stress response. (A) Heatmap of correlations of stress-induced gene expression among Bioprojects. Column labels indicate the Bioproject and reference. Row labels indicate the type of stress treatment. Datasets were hierarchically clustered based on correlation, and cell color indicates Pearson correlation. Columns are annotated based on inferred cluster assignment (A or B). Rows are annotated based on reported bleaching among stress-treated samples. (B) Confusion matrix showing holdout prediction frequencies for the validation set. Cells along the diagonal indicate correct predictions. (C) Density plot for control, cluster A stressed, and cluster B stressed samples along PC1 from Figure 2C. (D) Density plot for same samples along the DAPC axis from Figure 2D.

Functional enrichment in type A samples was consistent with the biological processes hypothesized to characterize the General Coral Stress Response. Previous studies have consistently linked several biological processes with coral stress, including downregulation of *cell division* and *ribosomes* and upregulation of *cell death*, *response to reactive oxygen species*, *NF-κB signaling, immune response, protein folding*, and *proteasomal protein degradation* (Barshis et al. 2013; Cziesielski et al. 2018; Aguilar et al. 2019; Cziesielski et al. 2019). We first examined differential expression values for all stressed type A samples compared to their controls. This analysis identified extensive functional enrichment consistent with the biological processes above (Table S5). We summarize these processes with representative GO terms selected from those most enriched in the type A response (Table S6).

In general, individual Bioprojects of type A showed consistent functional enrichment. To illustrate this, we repeated the enrichment analysis for each Bioproject individually and compared the results with those for all type A samples together (Figure 4A). In particular, *response to reactive oxygen* species (ROS), and *protein folding* were enriched for upregulation in all type A Bioprojects. One exception was ribosomes. Across the Bioprojects, *ribosome biogenesis* was sometimes upregulated and sometimes downregulated, with no enrichment for the type A samples as a whole (Figure 4A). One additional process, which we have summarized as *membrane vesicle* was also universally enriched in type A response.

**Figure 4:**
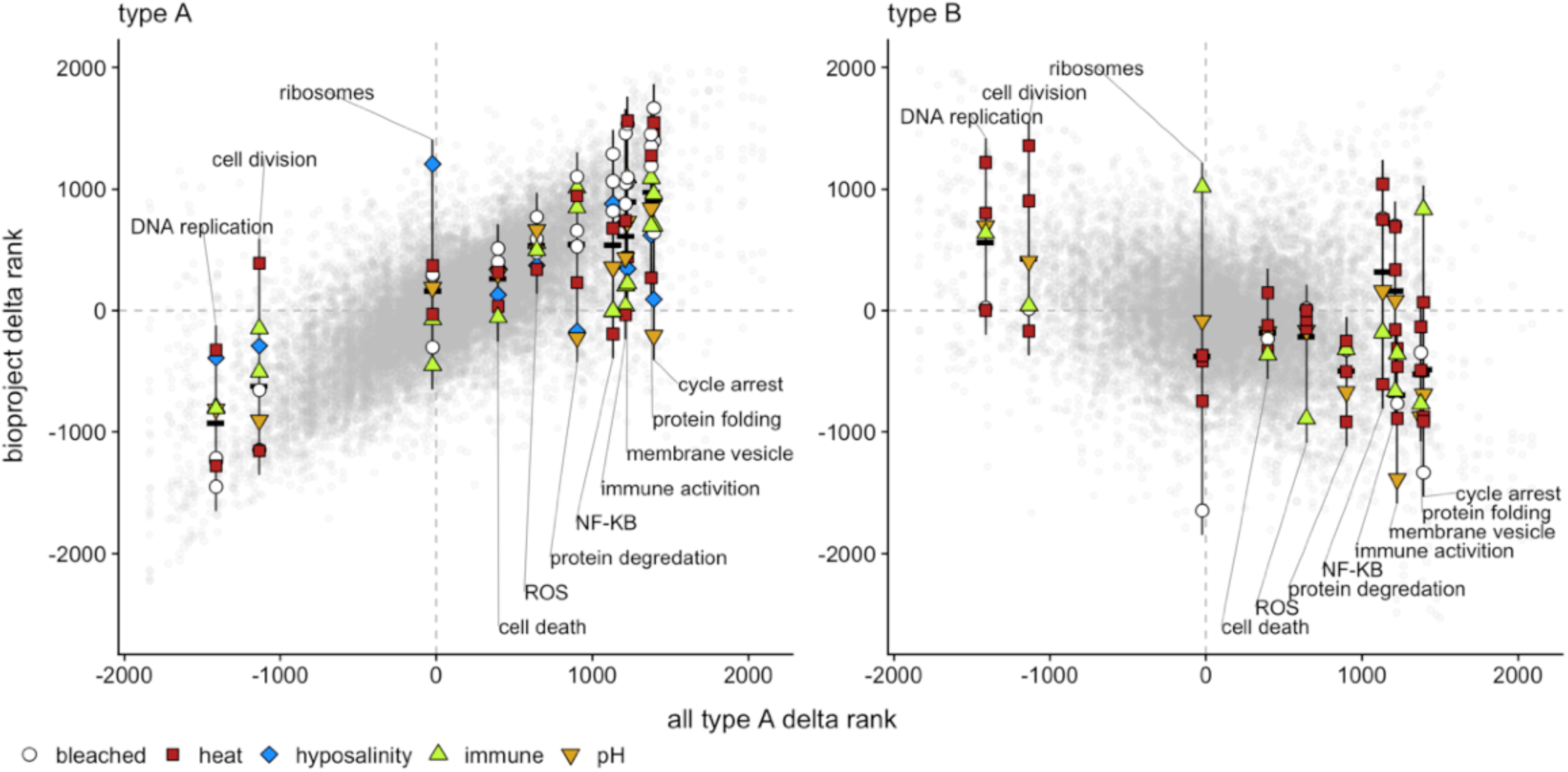
Gene ontology enrichment for selected terms hypothesized to characterize the general coral stress response. Delta ranks quantify the tendency of a GO term toward up-or downregulation in treated samples relative to controls. Negative values indicate preferential downregulation under stress treatment. Positive values indicate preferential upregulation. Each panel shows the relationship between delta ranks for individual Bioprojects and those based on all type A together. Each grey point represents a single GO term for a single Bioproject. Colored points connected by vertical lines show mean delta ranks for GO terms for the indicated biological process. These processes were selected because they have been previously hypothesized to characterize the general coral stress response. Point color indicates the type of stress used. Black horizontal dashes indicate the mean for each selected process. The left panel shows the type A Bioprojects. The right panel shows the type B Bioprojects. It is important to note that these plots are not intended to show statistical relationships, but are merely a convenient way of comparing functional enrichment across the different Bioprojects.

Functional enrichment for type B samples was highly distinct from cluster A. Here, terms linked with cell division tended to be upregulated relative to controls. Although variable, typically hypothesized stress-response processes tended to be downregulated (Figure 4B). Functional enrichment based on KOG class annotations further supported the distinction between the two response types. Enrichment in type A response largely reflected downregulation of cell division and metabolism genes, with upregulation of *Posttranslational modification, protein turnover, chaperones*, *Energy production and conversion*, *Translation, ribosomal structure and biogenesis*, *Signal transduction mechanisms*, and *Intracellular trafficking*, *secretion and vesicular transport* (Figure 5; Figure S7). As with the GO analysis, regulation of ribosomal genes was not consistent, sometimes tending toward upregulation, and sometimes downregulation. *Intracellular trafficking, secretion and vesicular transport* tended to be most strongly upregulated in heated and bleached samples (Figure S7). The type B response was highly distinct from A, and significantly negatively correlated (Figure 5). In particular, *Energy production and conversion* and *Translation, ribosomal structure and biogenesis* tended to be strongly downregulated in type B samples, whereas metabolism and cell division classes tended to be moderately upregulated. To further contrast the two stress responses, we repeated the KOG analysis on previously published gene expression data from yeast exposed to 19 different stressors. KOG enrichment for the yeast was more similar to the type B response, but did not resemble either of the coral responses particularly strongly (Figure 5). The yeast response and coral type B response were most similar (and distinct from type A response) in downregulation of *Translation, ribosomal structure and biogenesis*.

**Figure 5:**
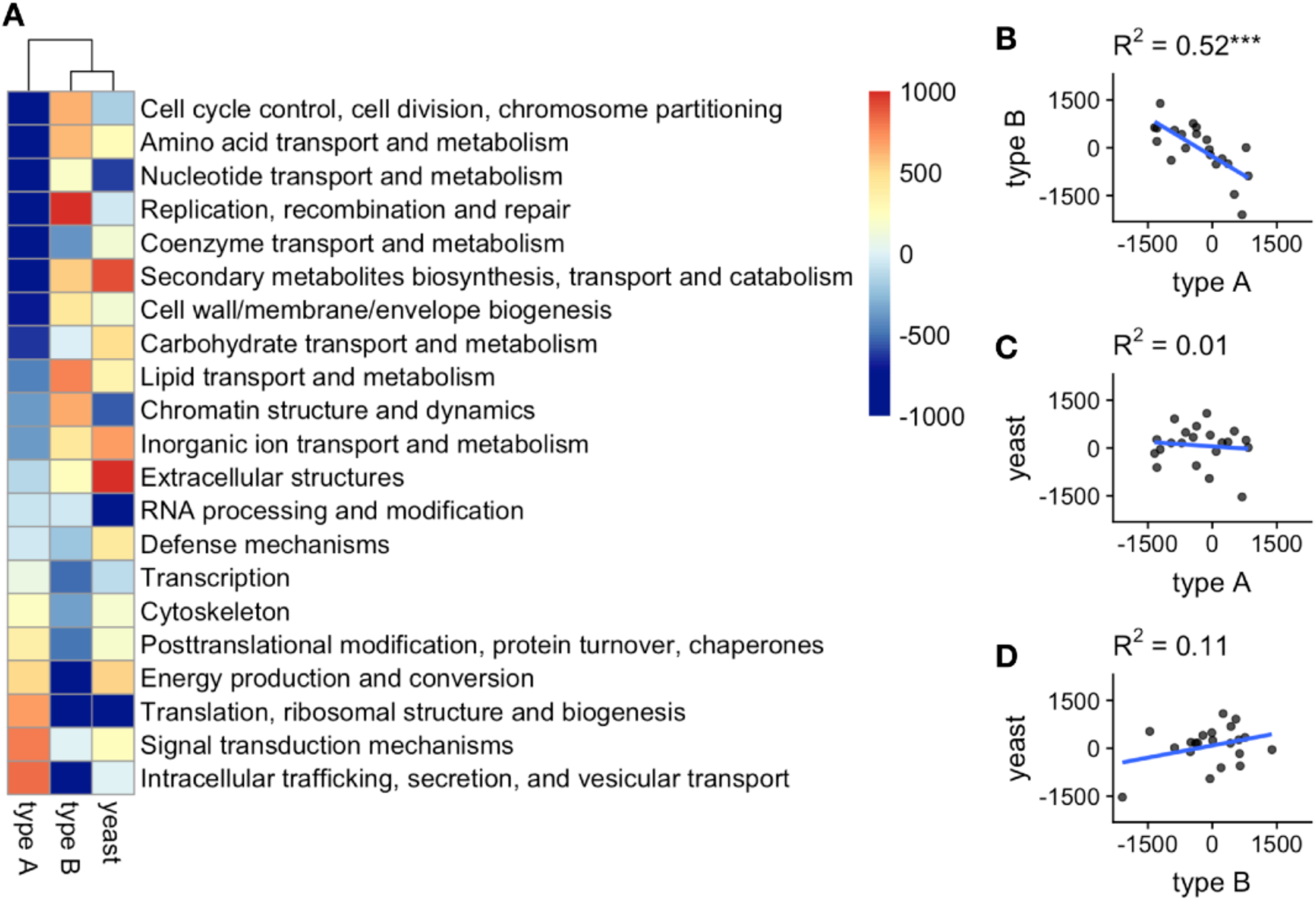
Summary of KOG class enrichment for the two stress types and for the yeast *Lachancea kluyveri*. (A) Heatmap of KOG class delta ranks, illustrating strength of enrichment for upregulation (red) or downregulation (blue) among KOG classes. (B-D) Scatterplots of relationship between KOG class delta ranks for different stress types.

A general stress response could be described in terms of a group of co-regulated genes that responds similarly to all types of stress. With this in mind, we used Weighted gene co-expression network analysis (WGCNA) to identify clusters (referred to as modules) of co-regulated genes and examined their association with stress treatments. The full WGCNA dendrogram and correlation heatmap are shown in (Figure S8). The three largest modules are shown in Figure 6. The red module appeared to capture the type A stress response. This module was upregulated for all types of stress in type A samples and enriched with gene ontology categories indicative of transcription factors, cell death, proteolysis, and growth inhibition (Figure 6; Table S7). The much larger green module, essentially capturing the background of the transcriptome (Figure S8), was downregulated by all types of stress among type A samples. The green module was functionally enriched with genes associated with growth and metabolism (Figure 5; Table S7). Notably, we did not detect a separate module for the type B response. Instead, type B samples appear to involve the same two modules that react to type A response (green and red) regulated in the opposite direction. This is consistent with our other results (Figure 3A, Figure 4B) and suggests that the type B response involves the same genes as type A but is inverted.

**Figure 6:**
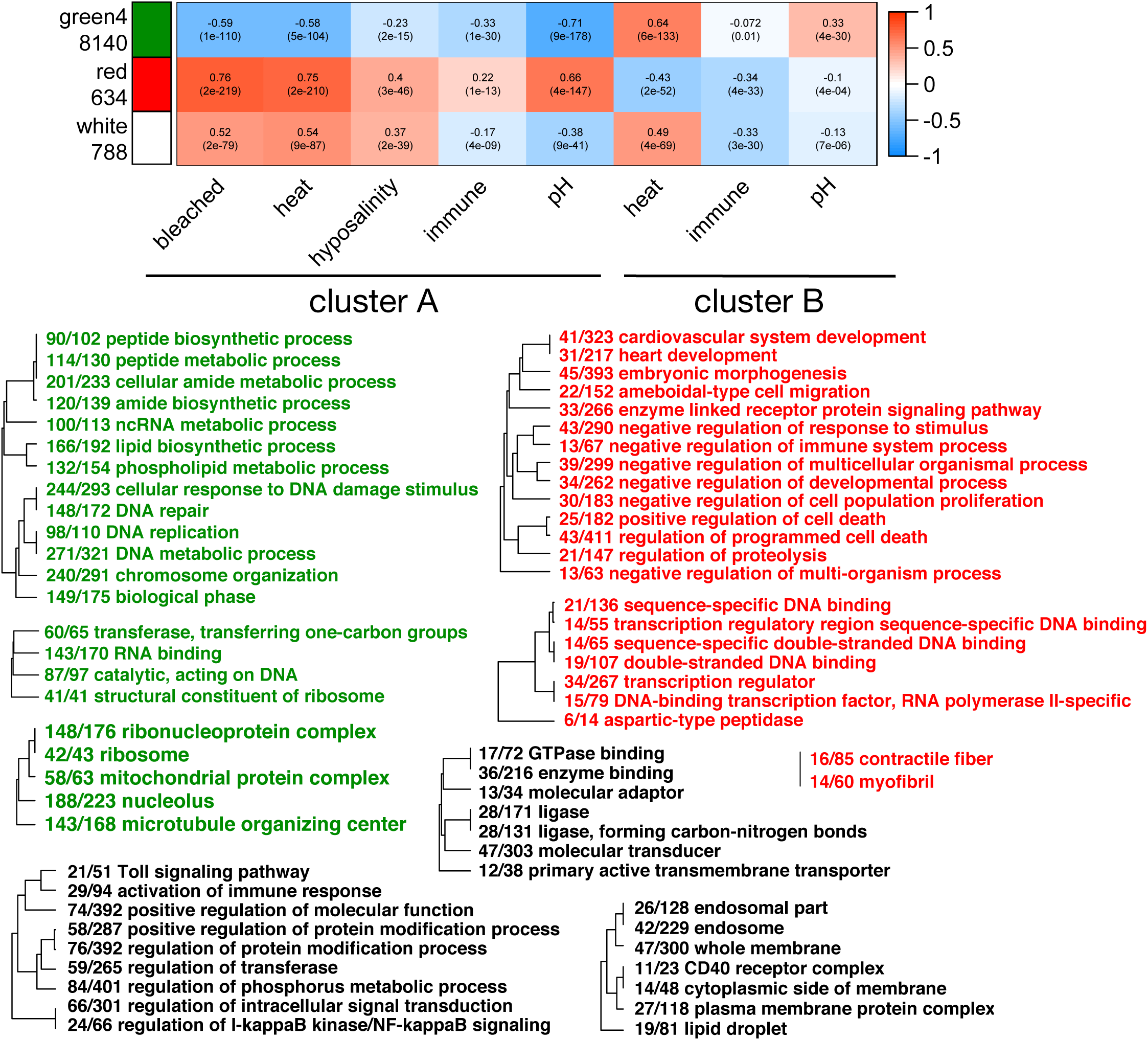
Sample trait correlations and gene ontology enrichment for three largest WGCNA modules. The heatmap illustrates correlations between the modules and stress treatments. The color scale indicates Pearson correlation. The number beneath each module title indicates the number of genes assigned to it. For correlation with stress treatments, samples were divided based on the clusters shown in Figure 3 (cluster A and cluster B). Color coded dendrograms show top enriched Gene Ontology terms for each module. All terms were enriched with a maximum FDR-adjusted p-value of 0.05 (Fisher’s exact test).

Expression of the white module was varied across stress types, upregulated by bleaching, heat, and hyposalinity, but downregulated by immune challenge and pH. Downregulation under immune challenge was especially surprising, as the module was enriched for genes associated with immune response as well as membrane vesicles and transporters (Figure 5; Table S7). The treatment associations and functional enrichment for this module suggest it may capture elements of the bleaching response. Indeed, in contrast to the green and red modules, membership in the white module was slightly more predictive of genes’ correlation with bleaching than with stress in general (Figure S9).

## Discussion

In this study, we analyzed publicly available and newly generated Tag-seq and RNA-seq data to assess the hypothesis of a general stress response in the genus *Acropora*. Contrary to expectations, we detected not one, but two classes of responses. The first of these, represented by cluster A in Figure 3, was consistent with the General Coral Stress Response. Responses among these Bioprojects tended to be correlated, and functionally enriched for previously hypothesized biological processes. Specifically, we detected significant upregulation for genes involved in *cell death*, *response to reactive oxygen species*, *NF-κB signaling, immune response, protein folding*, and *proteasomal protein degradation*. Surprisingly, the second cluster captured an entirely distinct transcriptional response, negatively related to type A.

WGCNA further supported these results. A module of 634 co-regulated genes and a contrasting module of over 8000 genes were up- and down-regulated by all types of stress for type A, but tended to be oppositely regulated in type B. Notably, the red module was enriched for factors inhibitory to cell proliferation (Figure 6). Although it remains unclear whether it operates in both directions, this negative regulatory relationship could potentially explain the paradoxically opposing responses observed, since growth-related processes tended toward upregulation in type B (Figure 4; Figure 5). A third module, enriched for immune, vesicle, and transport genes, was associated only with the types of stress used to induce bleaching. As this module was not induced by immune challenge (Figure 5), the enriched immune genes may involve interactions with symbionts. As the white module correlated with stress in projects that did not report bleaching (Figure 5; Figure S9), it may capture some of the regulatory responses that precede actual expulsion of symbionts.

The primary difference between the two response types appeared to be the severity of the stress. First, the median absolute log_2_ fold change between stress and control samples for type A was 1.3 fold that of type B (t-test p < 1e-9). This heightened response is suggestive of more severe stress in type A. This difference was further supported by the distribution of the stressed samples along the first principal component and the discriminant axis (Figure 3B-C). As these two axes largely captured transcriptomic variation between control and stressed corals, the tendency of type B samples toward their centers indicates less severe transcriptomic response, more similar to control samples. The frequency of bleaching was also much higher for type A (Figure 3A). Indeed, only one study in cluster B reported bleaching, and this was described as “mild coral bleaching” (Kaniewska, Chan, et al. 2015). Higher induction of bleaching matched with the generally higher temperatures used for heat treatment (median =34°C across type A Bioprojects; median = 32°C for type B). One exception was Parkinson et al. (2018), who used heat treatments of 35°C. As treatment duration in this study was only 1 hour, there may not have been sufficient time to mount the general response observed in type A. Among the three Bioprojects that applied immune challenges, the two in type A applied more severe stresses. Libro et al. (2013; PRJNA222758) sampled tissue that was at or adjacent to visible interfaces of white band disease, and Wright et al. (2017; PRJNA319662) physically abraded half their samples, and observed severe, albeit varied, mortality in the immune challenged group. In type B on the other hand, Weiss et al. (2013; PRJNA200542) injected their samples with immunogens: muramyl dipeptide and polyinosinic:polycytidylic acid, which were intended to elicit a specific immune response by mimicking infectious agents. In general, pH appeared to elicit less severe expression changes than other treatments. Indeed, reports of the physiological severity of elevated pCO_2_ on corals are varied (Langdon and Atkinson 2005; Rodolfo-Metalpa et al. 2010), and corals may very well be capable of resilience (McCulloch et al. 2012). The only study using pH treatment to show the type A response was Moya et al. (2015; PRJNA260269), whereas a similar experiment (Moya et al. 2012; PRJNA149513) was type B. The difference here was a prolonged exposure (9 days beginning immediately post-fertilization in Moya et al. 2014) rather than short term (3 days beginning immediately after settlement in Moya et al. 2012).

To summarize, the main difference between the two stress types appeared to be severity of stress induced, with higher stress in type A. This is consistent with the proposed role of the yeast ESR, to serve as a blanket response that prepares cells to protect themselves under diverse types of stress (Gasch et al. 2000; Gasch 2003). The weak correlations observed within type B may instead reflect specific transcriptional responses to different stressful conditions that did not reach levels sufficient to induce the General Coral Stress Response. Hence contrary to expectations, the severity of a stress treatment may not only affect the magnitude of transcriptional response but its functional nature. This fact could have critical importance when predicting physiological responses based on comparative transcriptomics.

The surprising negative relationship between the two response types may explain previous inconsistencies between gene expression studies. For instance, Barshis et al. (2013) hypothesized that corals adapt to stressful conditions by constitutively expressing stress response genes at higher levels (transcriptional “frontloading”). According to this hypothesis, constitutive profiles more similar to stressed individuals should indicate higher tolerance. Contrary to this hypothesis, Dixon et al. (2015) found that larval cultures with greater transcriptomic similarity to heat-stressed adults had lower heat tolerance. Here we find that the adult heat treatment from Dixon et al. (2015)(31.5°C for 72 hours) fell into the type B response, which is opposite the type A response observed under more severe heat stress treatments, such as that applied in Barshis et al. (2013). In other words, if Dixon et al. (2015) had stressed their adult corals more severely, their results may have supported “frontloading” *sensu* Barshis et al. (2013).

Gene annotation is a major challenge in ecological genomics. For instance, with this dataset, we identified a final set of 25 genes that were members of the red module and upregulated for every type A Bioproject with a minimum fold change of 1.5 (log_2_ fold change > 0.59)(Table S8). Among the annotated genes in this set were two were heat shock proteins, two oxidase/oxidoreductases, two involving retro-transposition, a TNF receptor-associated factor, a matrix metallopeptidase, and a transcription factor. Thirteen of these genes (52%) however, lacked annotations. Given the diversity of studies included, these genes can be confidently labeled as stress response genes. Based on our results we have generated contextual annotations for *A. millepora* genes including bleaching (|log_2_ fold change| > 0.59 in all type A bleached Bioprojects) and two tiers of general stress response (the core set described above and a more lenient set with |log_2_ fold change| > 0.32 in >80% of type A Bioprojects)(Table S8). Additional sets with greater or lesser stringency could be generated as preferred (Table S9; Table S10). We hope that this first effort will improve in scope and detail by adding more gene expression studies in the future.

## Supporting information

Supplemental table S1: original SRA table

Supplemental table S2: sample traits

Supplemental table S3: read counts

Supplemental table S4: eggNOGmapper annotations

Supplemental table S5: all type A GO_MWU results

Supplemental table S6: custom GO sets

Supplemental table S7: WGCNA module GO_MWU results

Supplemental table S8: contextual annotations

Supplemental table S9: Bioproject DESeq results

Supplemental table S10: WGCNA module membership

## Data availability

Reads generated for this study have been uploaded to the SRA database project accession PRJNA559404. Scripts for data processing and analysis, as well as intermediate datasets are available on Github: https://github.com/grovesdixon/Acropora_gene_expression_meta.

---

**Figure S1:**
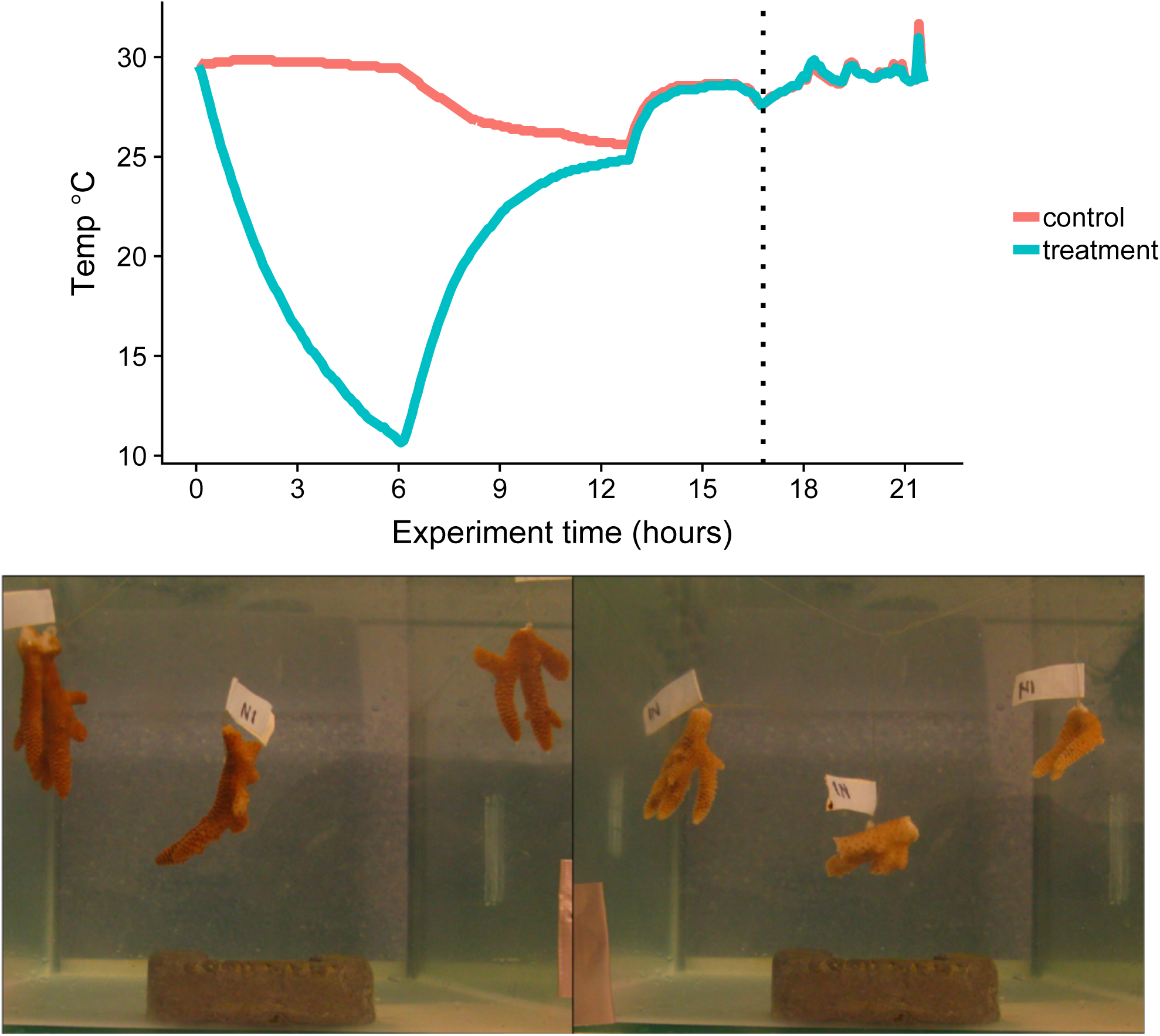
Temperature trace for the cold experiment. Each treatment was conducted in a 10 L container each with three replicates from each of the four colonies (12 nubbins per treatment group). The cold treatment container was moved to a refrigerator set to 4°C for six hours while the control container was placed in a cabinet to mimic the darkness experienced in the refrigerator. After six hours both containers were returned to outdoor tabletops and air circulation was resumed. After six more hours flow of filtered seawater was resumed. Tissue samples were fixed in ethanol 17 hours after starting the treatments (indicated by black dotted line) and immediately placed at −80 °C. Example photos show control (left) and treated (right) samples.

**Figure S2:**
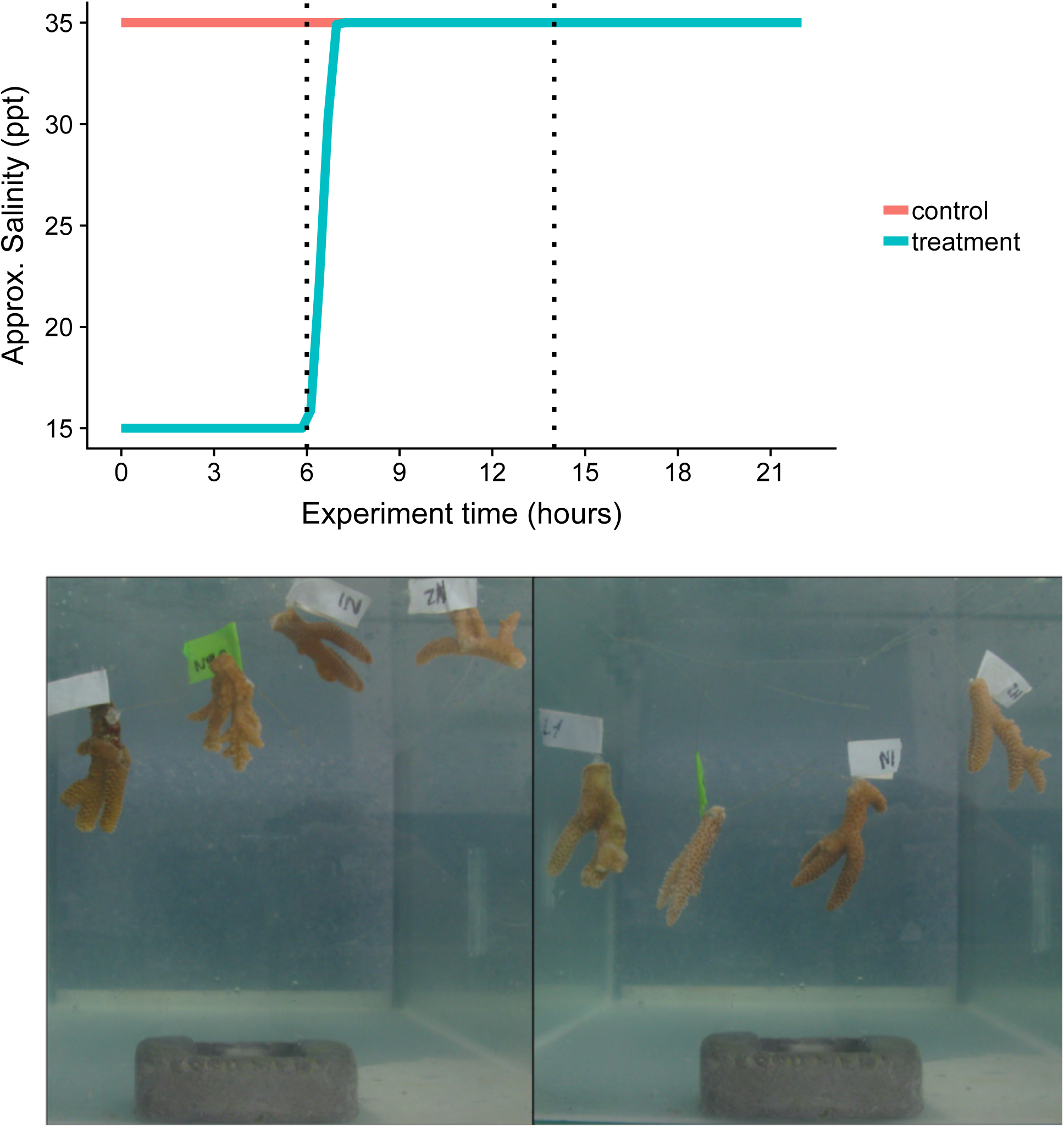
Trace of approximate salinity for hyposalinity experiment. Each treatment was conducted in three 5 L containers per treatment group, each with a single replicate nubbin from each colony (12 nubbins per treatment group). The hyposalinity treatment began with immediate submersion in 42.8% filtered seawater (approximately 15 ppt), maintained with air circulation for six hours of exposure before resuming flow of filtered seawater. Tissue samples were fixed in ethanol at two time points: 6 and 14 hours after starting the treatments (indicated by black dotted lines). Fixed samples were stored immediately placed at −80 °C. It should be noted that salinity was not measured directly, so the traces represent an approximate estimation. Example photos show control (left) and treated (right) samples.

**Figure S3:**
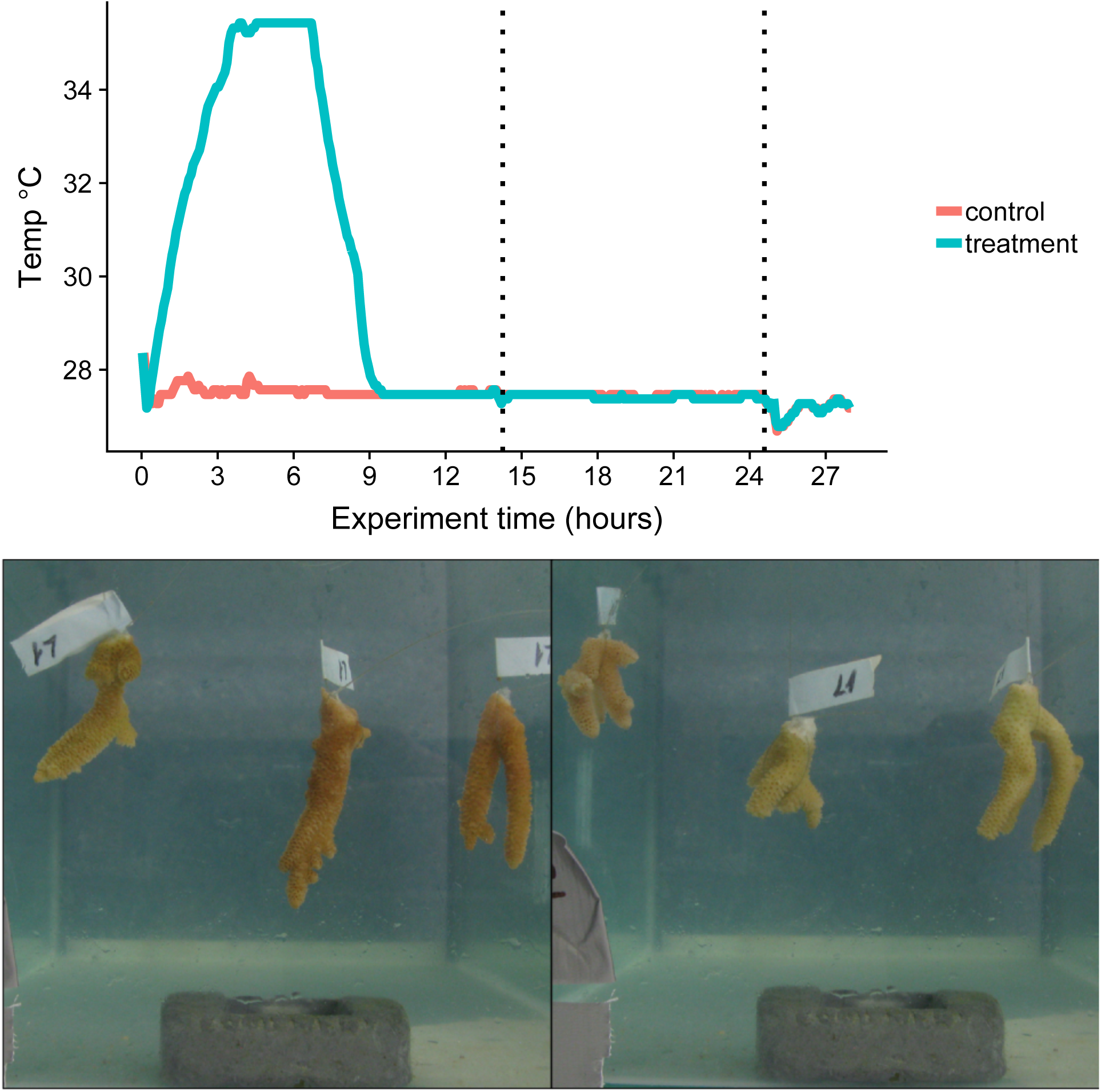
Temperature trace for first heat experiment. Each treatment was conducted in a 10 L container each with three replicates from each of the four colonies (12 nubbins per treatment group). The heat treatment container was ramped to 36°C over three hours and held at 36°C for three additional hours. After six hours both containers were returned to outdoor tabletops and air circulation was resumed, ramping the treated group back to 28°C over three hours. After six more hours flow of filter seawater was resumed. Tissue samples were fixed in ethanol at two time points: 14 and 25 hours after starting the treatments (indicated by black dotted lines). Fixed samples were stored immediately at −80 °C. Example photos show control (left) and treated (right) samples.

**Figure S4:**
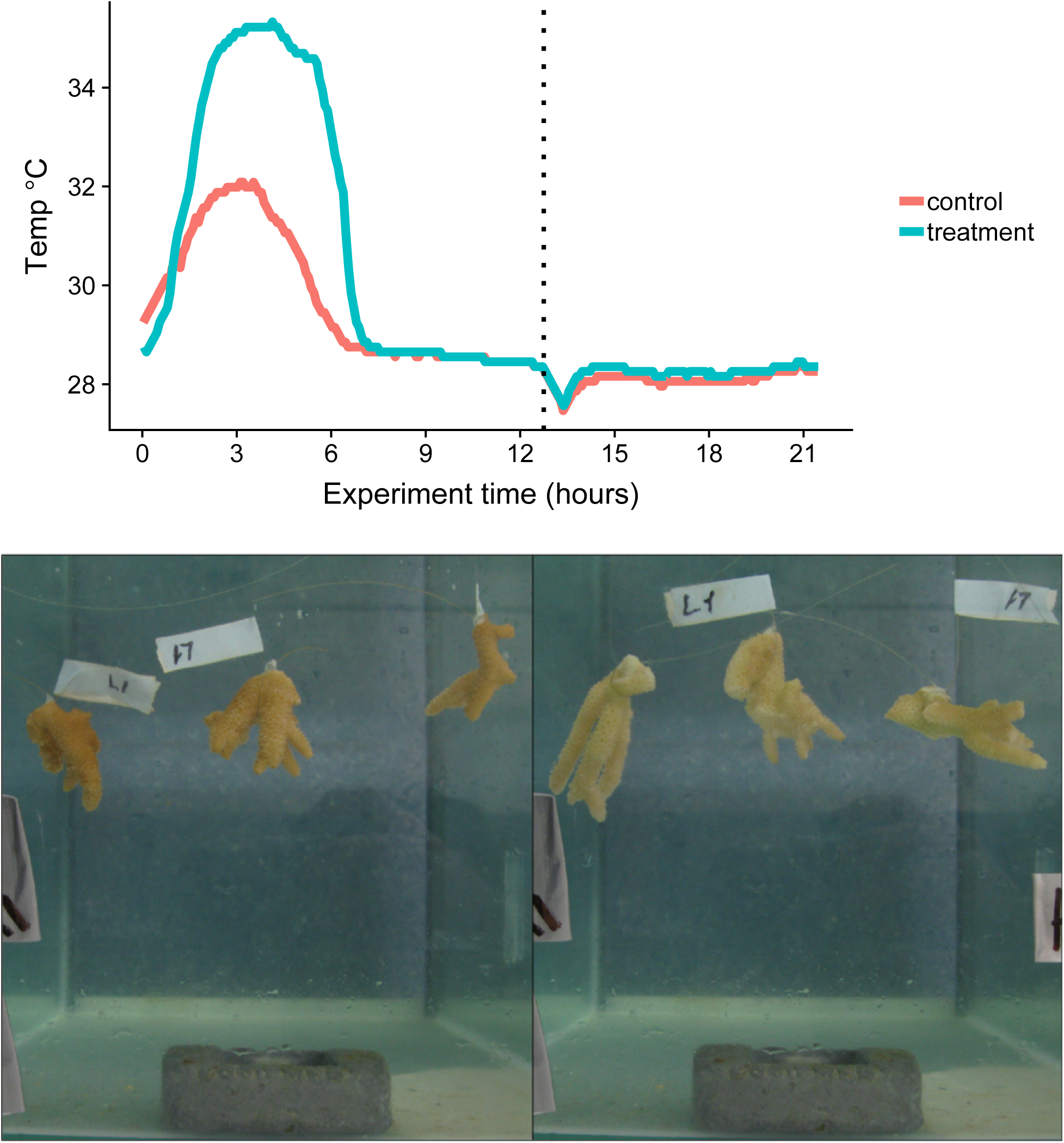
Temperature trace for the second heat experiment. Each treatment was conducted in a 10 L container as part of the multi-stress experiment, each with three replicates from each of the four colonies (12 nubbins per treatment group). The heat treatment container was ramped to 35°C over three hours and held at 35°C for three additional hours before ramping back to 28°C over three hours. After six hours the heat and control containers were returned to outdoor tabletops and air circulation was resumed. Tissue samples were fixed in ethanol 13 hours after the experiment began. Fixed samples were stored immediately at −80 °C. Example photos show control (left) and treated (right) samples.

**Figure S5:**
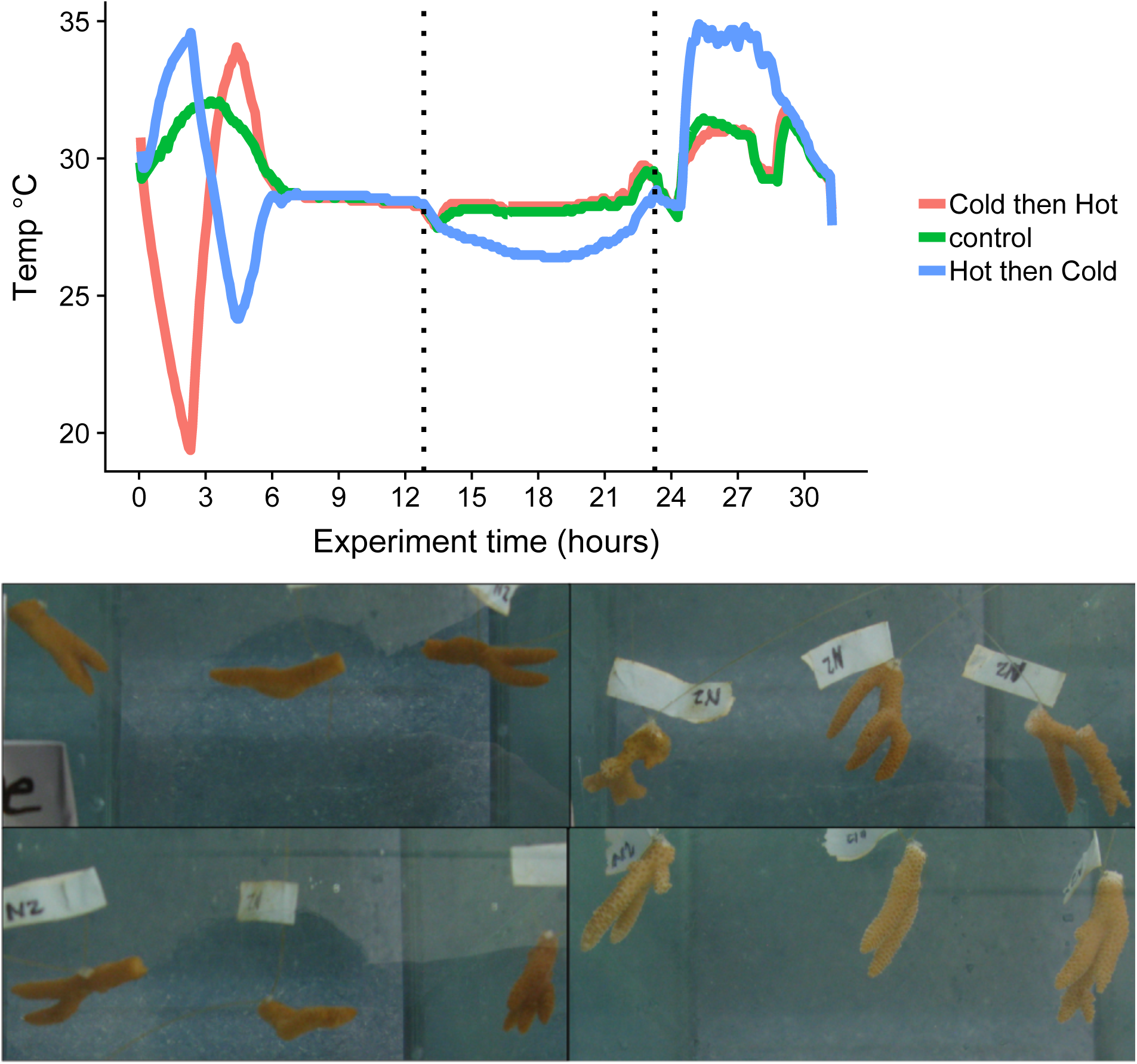
Temperature traces for multi-stress experiment. Each treatment was conducted in a 10 L container each with three replicates from each of the four colonies (12 nubbins per treatment group). The hot-then-cold treatment group was ramped to 35°C over three hours, and then moved to a refrigerator at 4°C for three hours, then returned to an outdoor tabletop. The cold-then-hot treatment group was first placed in the refrigerator at 4°C for three hours, then ramped to 35°C over three hours before it was moved to an outdoor tabletop. Tissue samples were fixed in ethanol at two time points: 13 and 23 hours after starting the treatments (indicated by black dotted lines). Fixed samples were stored immediately at −80 °C. Example photos show control (left) and treated (right) samples: cold-then-hot (top), hot-then-cold (bottom).

**Figure S6:**
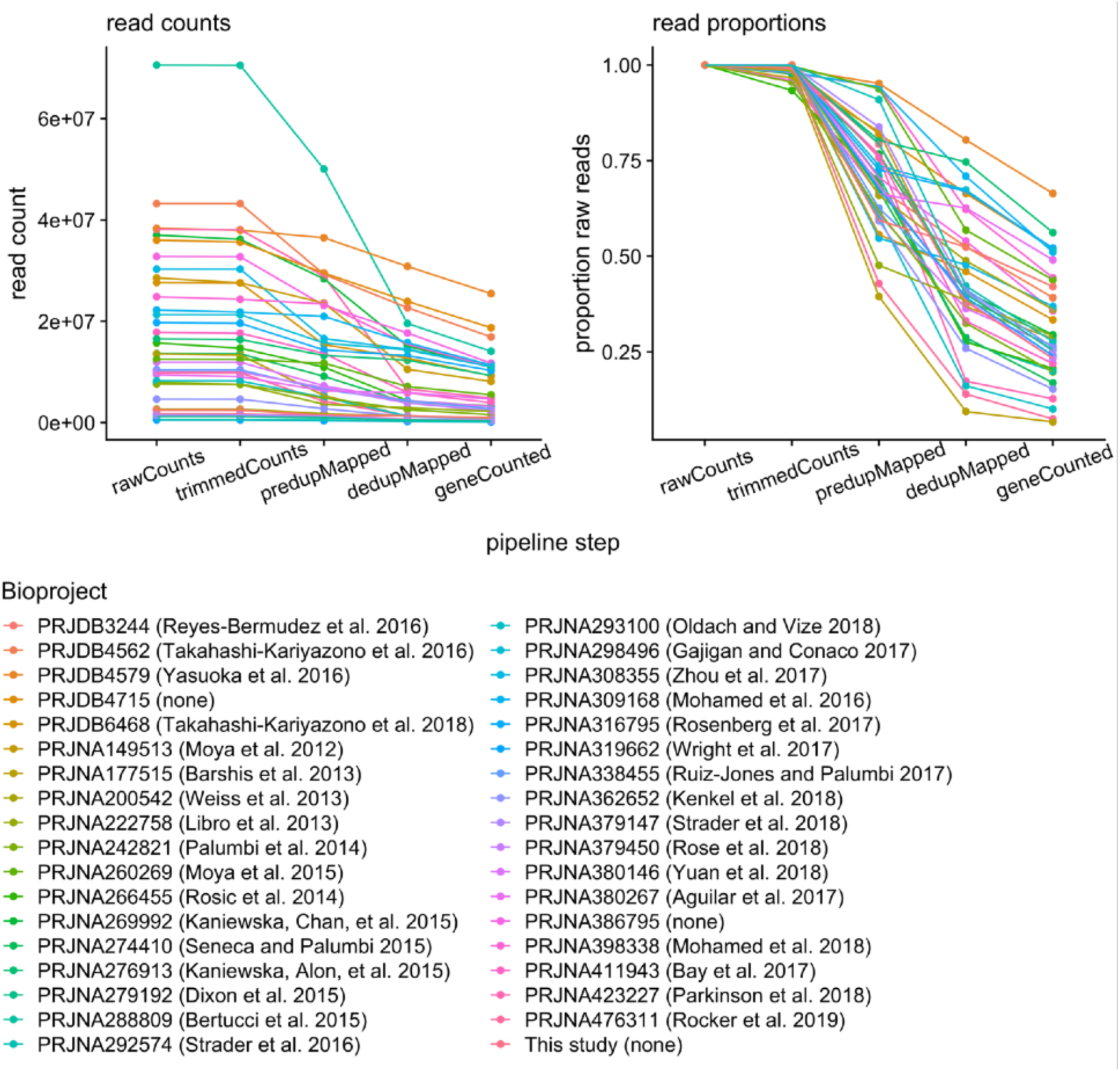
Mean read counts per sample for each Bioproject throughout data processing pipeline. Absolute read counts (left) and proportion of starting read counts (right).

**Figure S7:**
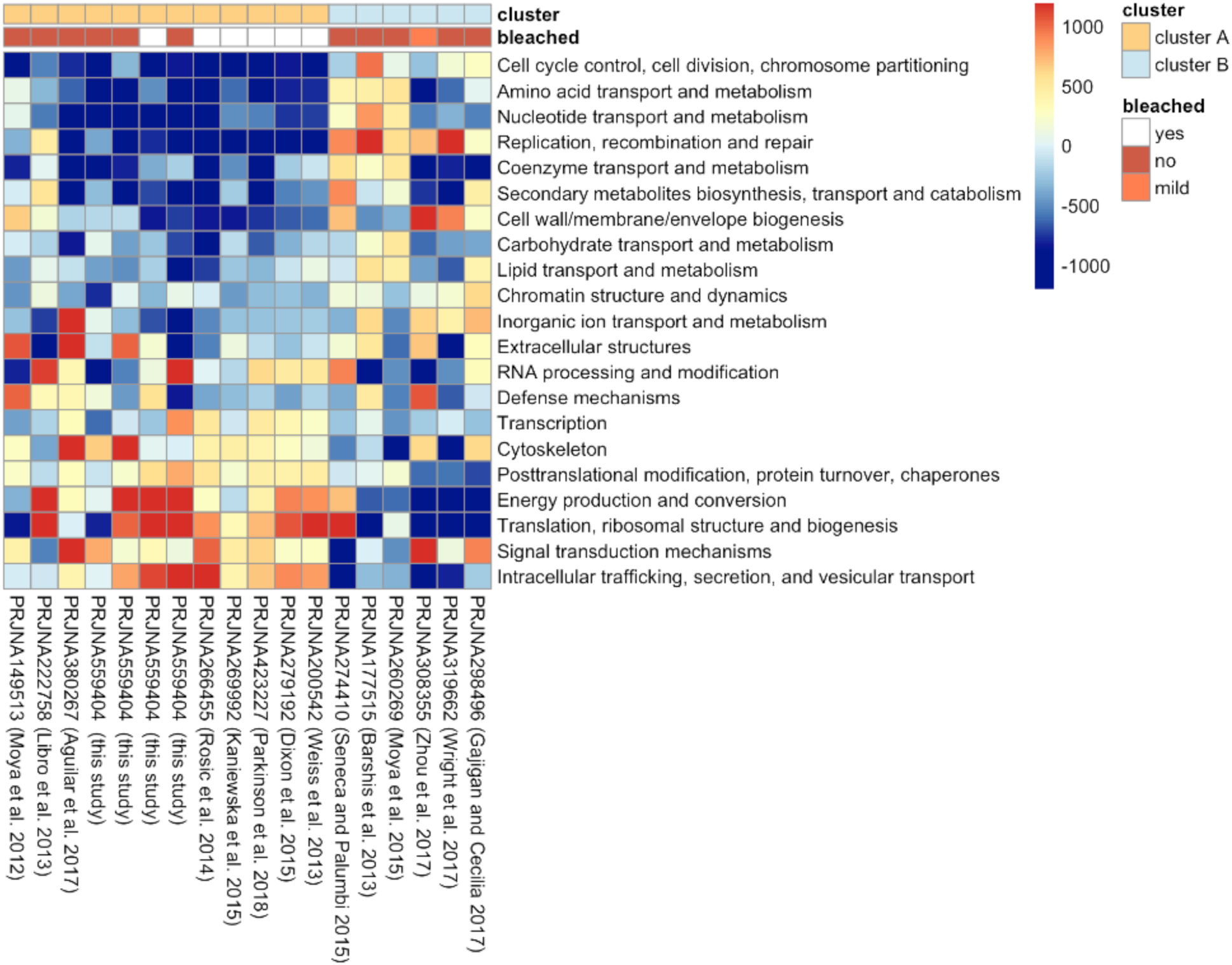
Heatmap of delta ranks quantifying tendency of KOG classes toward up- (red) or down regulation (blue) in stress treated samples. Each row represents a KOG class, each column represents a Bioproject. Ordering of Bioprojects is set to match the clustering in Figure 3.

**Figure S8:**
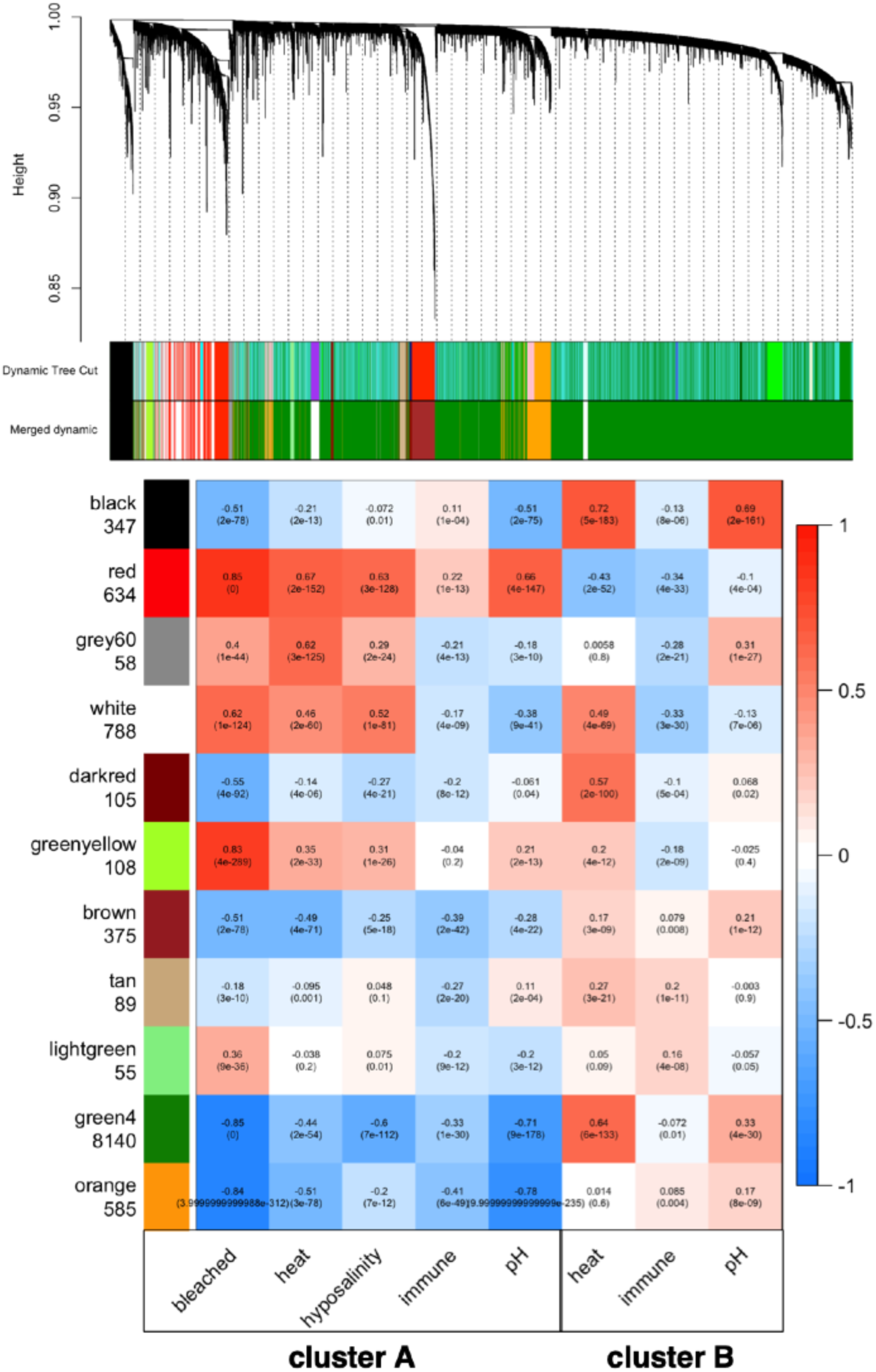
Module dendrogram and sample trait correlation heatmap from WGCNA. Vertical lines in dendrogram indicate individual genes. Module color assignments are shown beneath. Dynamic Tree Cut indicates the module assignments based on a minimum module size of 30. Merged dynamic indicates the modules after merging at a selected threshold of 0.3. Heatmap illustrates correlation between module eigengenes and sample traits. Within each cell, the Pearson correlation is given above and the p-value below.

**Figure S9:**
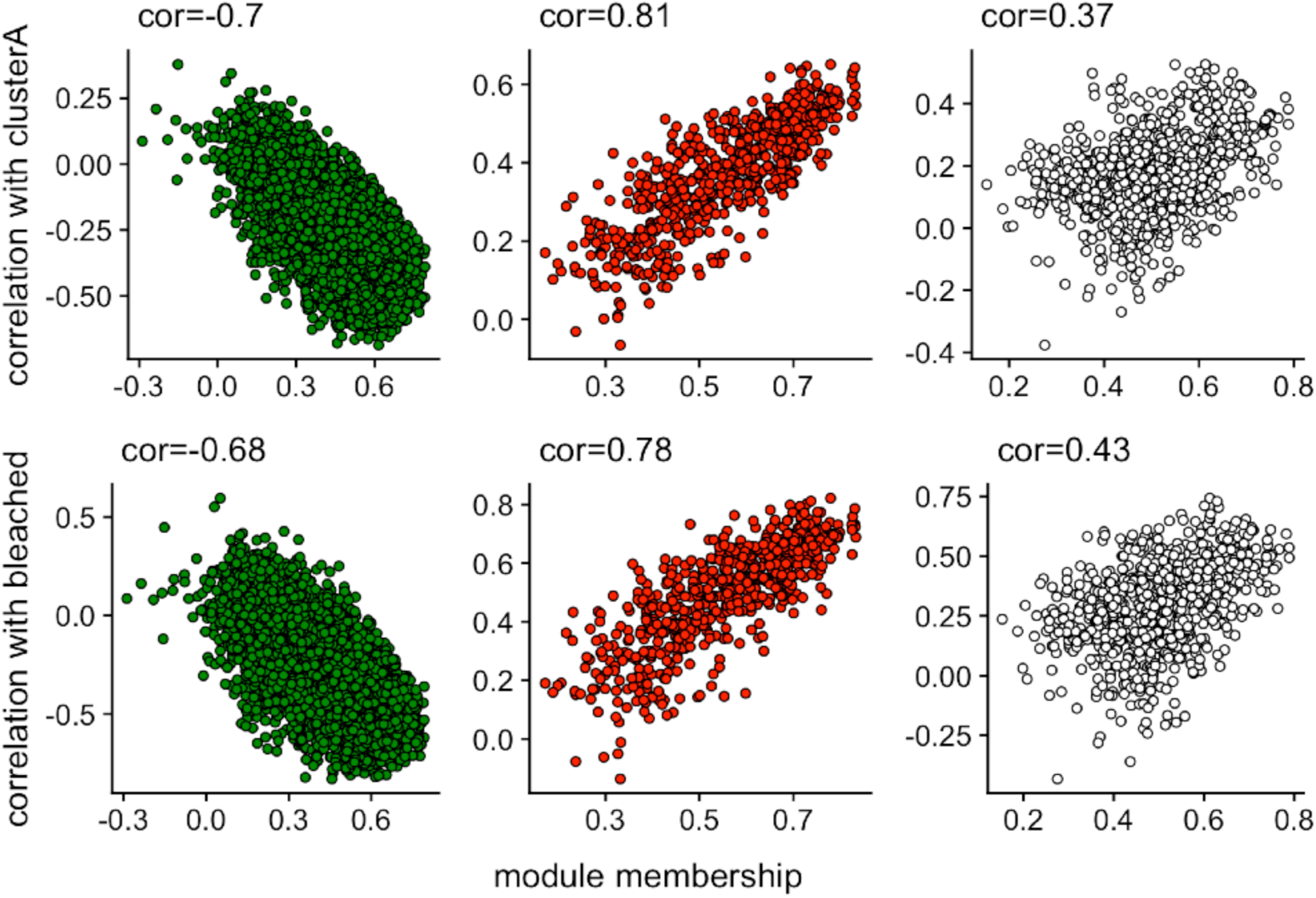
Scatterplots of module membership and trait correlation for the three largest modules. Each panel shows genes’ module membership values on the X-axis (the correlation of the genes’ expression levels and the module eigengene), and the genes’ correlation with the indicated trait on the Y-axis (top panel = stress treatment in type A samples; bottom panel = bleaching in type A samples). For all three modules, the relationship is in the same direction, indicating the membership in each module predicts the same response due to bleaching and stress in general. Genes’ correlations with bleaching tend to be stronger than with stress in general (compare scales of Y axes). For the red and green modules, the relationship between module membership and trait correlation is stronger for stress in general. In contrast, for the white module, it is stronger for bleaching. Hence, membership in the white module predicts genes’ correlation with bleaching slightly better than for stress in general.

